# Minimal diadenylate cyclases have been co-opted to detect phage immune evasion

**DOI:** 10.64898/2026.08.10.743972

**Authors:** Ali Nabhani, Ashley E. Sullivan, Naryung Oh, Giulia F. Otsuki, Laurel K. Robbins, Jeslyn K-Y. Lee, Daniel S. Izrailevsky, Charlotte R.K. Hoffman, Aaron T. Whiteley, Benjamin R. Morehouse

**Author notes:** equal contribution (Ali Nabhani, Ashley E. Sullivan).

## Abstract

Many bacterial immune defenses transmit recognition of phage infection via the generation of diverse cyclic nucleotide second messengers. Phage have evolved to subvert this kind of immunity by sequestering, degrading, or inhibiting synthesis of these signaling molecules. Consequently, bacteria have evolved counter-mechanisms to detect disruption of cyclic nucleotide signaling and induce another layer of immune protection. Here we detail our discovery of the PanDA defense system (<u>P</u>anoptes-like <u>D</u>is<u>A</u>), an antiphage defense which detects sequestration of 3′3′-c-di-AMP by phage sponge proteins. PanDA consists of two proteins, PanS and PanE, which are both necessary for defense. PanS contains a minimal diadenylate cyclase (DAC) domain that constitutively generates the cyclic dinucleotide 3′3′-c-di-AMP which binds to and represses a toxic effector, the 2TM-β family protein PanE. When a cell is infected by a phage encoding the sponge protein Acb4 (anti-CBASS protein 4), PanE is activated and induces membrane permeability. This work represents the first confirmed use of 3′3′-c-di-AMP as an immune second messenger in bacteria, facilitated by the exaptation of a DAC domain which has thus far only been best understood for its non-immune signaling roles.

## INTRODUCTION

Bacteria face a prolific number of selection pressures in their native environments. Foremost among these pressures, predation upon bacteria by bacteriophage (phage) has selected for the rapid evolution and diversification of systems to defend against infection^1^. Such bacterial immune systems make use of a wide variety of mechanisms to recognize infection, signal its occurrence, and prevent phage replication. Certain defenses target and degrade phage nucleic acids in a sequence specific manner, including the well-known restriction modification and CRISPR antiphage systems. Others, such as the Hachiman defense system, degrade nucleic acids non-specifically upon protein-DNA interactions^2^. Alternative systems such as bacterial gasdermins, Thoeris, Pycsar, and CBASS, rely on protein-protein interactions to recognize phage and induce defense via membrane impairment, metabolite depletion, and a variety of other effector functions^3–6^.

In turn, phage have evolved a plethora of anti-defense proteins to subvert bacterial immune systems^7^. Mechanisms of phage anti-defense include direct inhibition of defense system components via binding or covalent modification, modification of phage nucleic acids to avoid detection, synthesis of metabolites depleted by defense systems, and the sequestering or degradation of bacterial immune second messengers^7,8^. The recently discovered Panoptes system calls attention to yet another layer in the evolutionary arms race between bacteria and phage in which bacteria have evolved to recognize phage perturbation of cyclic nucleotide-based signaling^9,10^. The use of cyclic nucleotides as second messengers to propagate bacterial immune signals is a reoccurring theme which bridges several of the systems mentioned previously such as CBASS, Pycsar, Thoeris, and Panoptes^11^. This theme extends beyond bacterial immunity as components of CBASS and Thoeris systems are conserved in organisms as distantly related as humans in the form of the cGAS-STING and TLR-related pathways^12–15^.

However, cyclic nucleotides, particularly mono- and dinucleotides, are also critical components of core processes outside of immunity across the tree of life. In bacteria and *S. cerevisiae*, cAMP serves several roles including as a regulator of carbon-source operons, growth rate, and resistance to various stress factors^16,17^. In humans, cAMP and cGMP have roles including, but not limited to, metabolic regulation, muscle contraction, phototransduction, and neurotransmitter modulation. The cyclic dinucleotides 3′3’-cyclic-di-GMP (c-di-GMP) and 3′3’-cyclic-di-AMP (c-di-AMP) have well established roles in osmotic/potassium homeostasis and biofilm formation, respectively^18,19^. Fewer roles of cyclic dinucleotides in eukaryotes have been discovered, though it has been found that c-di-GMP triggers stalk cell differentiation in *Dictyostelium*^20^.

The use of cyclic nucleotides in both immune and core processes across such diverse taxa poses several evolutionary questions. Much attention has been paid to the evolution of subsets of prokaryotic immune domains into the immune and core machineries of organisms as distantly related as humans^15,21,22^. By comparison, much less attention has been paid to the exaptation of core prokaryotic domains for use in bacterial immunity. c-di-AMP and structural isomers have been characterized as the primary products of cyclase homologs belonging to the Panoptes system while other isomers likely made by uncharacterized homologs of the same or other systems^9,10^. Additionally, new findings support that archaeal TIR-SAVED effectors demonstrate an ability to sense host-derived c-di-AMP when expressed in a non-endogenous environment^23^.

The diadenylate cyclase (DAC) domain is the primary producer of c-di-AMP and is well characterized as the crux of many regulatory processes across a diverse range of bacteria^24,25^. DACs have recently been bioinformatically noted for their frequent association with certain CBASS systems^26^. One such DAC-containing system in *E. coli* provided defense against phage infection in a catalytic-dependent manner, though an in depth interrogation of the signaling mechanism behind defense has not been performed^27,28^. Here we use a combination of bioinformatics, microbiology, structural biology, and biochemistry approaches to characterize a novel Panoptes-like system that incorporates a minimal DAC. We characterize the phylogenetic and sequence conservation of this novel minimal DAC and show that it uses c-di-AMP signaling to respond to phage disturbance of the cellular cyclic nucleotide pool.

## RESULTS

### A clade of predicted immune operons incorporates a minimal DAC domain-containing protein

Prokaryotic DACs were recently organized into 22 proposed families based on their regulatory domain organization^26^. To further chart the diversity of bacterial DACs, we searched ∼250,000 publicly available bacterial proteomes, seeding with members of the proposed families alongside sequences of several published DAC homologs. Our search yielded approximately 1200 non-redundant hits which were aligned with MAFFT and used to construct a maximum likelihood tree (**Figure 1a**). In terms of taxonomy, the species represented on the tree belonged to a wide range of phyla from several distinct evolutionary niches, including both gram-positive and gram-negative organisms.

**Fig. 1:**
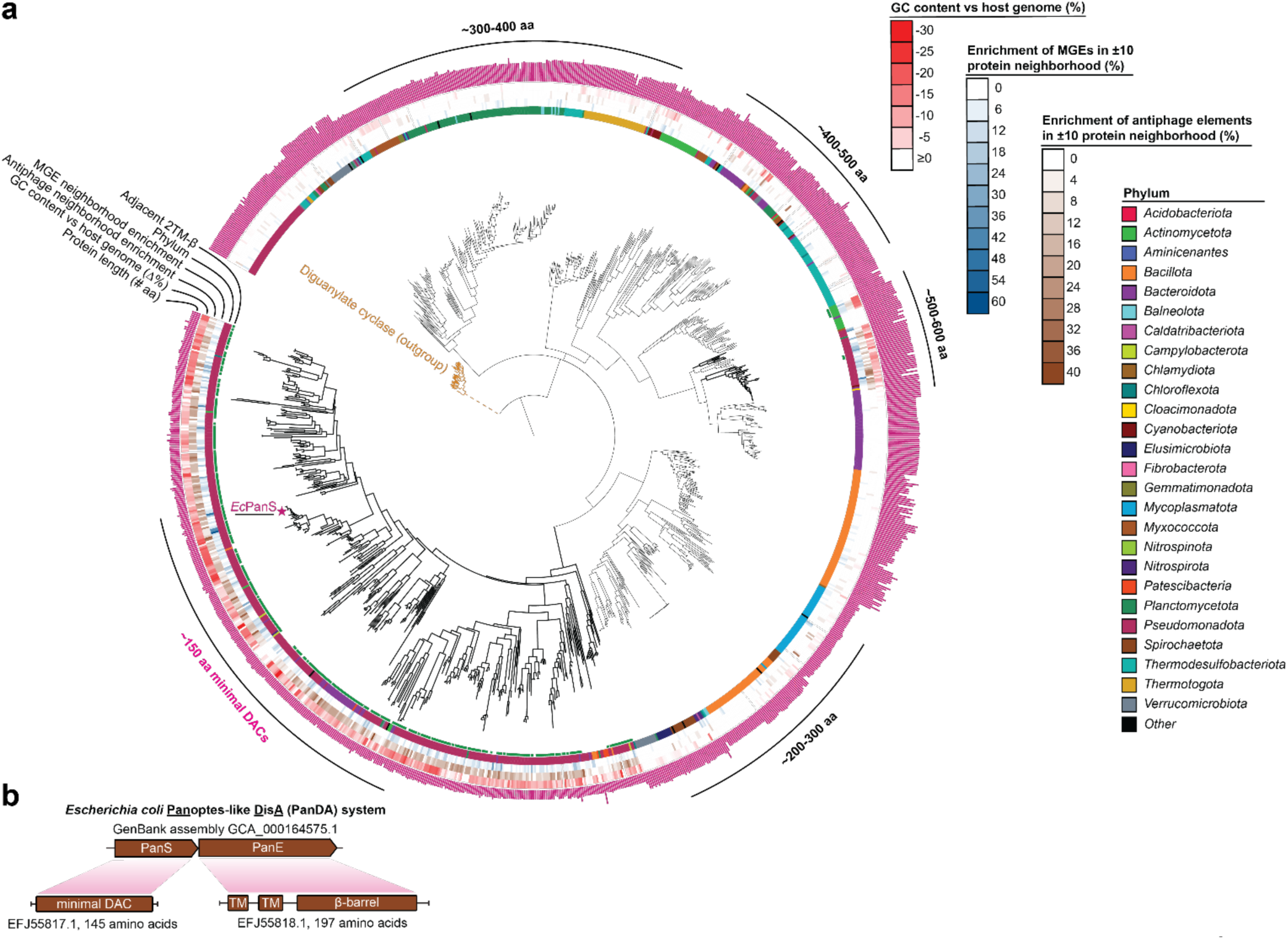
A family of minimal DAC domain-containing proteins are implicated in antiphage immunity. a) Phylogenetic tree of DACs collected during the initial search performed in this study. The presence of an adjacent 2TM-â effector is denoted by a dark green box on the innermost ring. Phyla from which the DACs on the tree originate are denoted by the second ring from the center of the tree. MGE enrichment is represented in the third ring where dark blue indicates a greater number of MGEs in the protein neighborhood. Antiphage enrichment is represented in the fourth ring where dark brown indicates a greater number of antiphage elements in the protein neighborhood. The difference in GC content between the coding sequence for each hit and the host genome is denoted by the red heatmap in the fifth ring. Protein length in amino acids is denoted by the outermost ring of pink bars surrounding the tree. The diguanylate cyclase outgroup used here is colored gold. b) Operon and domain organization of an *E. coli* PanDA system which was selected for further study.

Next, we graphed the length of each hit in amino acids and observed a clustering of DACs which generally coincided with protein length (**Figure 1a**). Predicted structural analysis revealed a diversity of domain organizations, several of which have yet to be described in the literature (**Extended Data Figure 1**). Interestingly, hits within a clade primarily belonging to the phylum *Pseudomonadota* were only ∼150 amino acids in length (**Figure 1a**). Such a short length corresponds solely to the DAC, lacking any of the appended regulatory domains which characterize other homologs. Indeed, AlphaFold3 models indicated that these sequences resembled a minimal form of a protein solely composed of the DAC domain (minimal DAC, mDAC). The lack of a regulatory domain also implied an alternative means of activity regulation or a complete lack thereof.

To identify DACs across our tree which might be involved in antiphage defense, we examined the protein neighborhood (±10) of each hit for indications of immunity. Bacterial defense genes have been frequently identified by their colocalization within “defense islands”, genomic groupings of antiphage genes which can be more easily transferred between bacteria and populations^29,30^. We quantified the proportion of the protein neighborhood of each hit which contained predicted antiphage proteins and found that members of the mDAC clade were significantly enriched in predicted antiphage neighbors (**Figure 1a**). Two additional branches composed of hits ∼575 amino acids in length were similarly enriched in antiphage neighbors, with structural analysis suggesting that they belonged to the recently formed CdaG family of DACs.

Antiphage systems are often encoded within mobile genetic elements (MGEs) such as prophages, integrons, and plasmids due to the fitness benefit which they provide their respective bacterial host^31–34^. As such, we also quantified the presence of MGE-related proteins within each hit’s genomic neighborhood and found a significant, but less uniform, enrichment of MGEs in the vicinity of the same clades/branches of DACs across the tree. A difference in GC content compared to the host genome is a telltale sign that a genetic element has been transferred between bacterial genomes^35^. Further, it has been noted that defense genes often have GC contents lower than their host genome^28,36^. Building on the hypothesis that the mDAC clade and two branches of longer DACs may be involved in phage defense, we calculated the difference in GC content between each coding sequence represented on our tree and the respective host genome (Figure1A). We found an average difference in GC content of ∼10% across these groups, further indicating that these genes may be involved in defense.

To gauge if any of our DACs belonged to recognizable defense systems, we examined the proteins directly adjacent to each hit. Surprisingly, greater than 90% of the mDAC clade appeared operonic with predicted effectors composed of two transmembrane helices appended to a β-barrel (2TM-β). The presence of a minimal cyclase (mDAC) operonic with a 2TM-β resembled a family of the recently discovered Panoptes system^9,10^. We therefore named this putative defense operon the **<u>Pan</u>**optes-like **<u>D</u>**is**<u>A</u>**system (PanDA) in reference to the first discovered DAC, the DisA protein (**Figure1b**)^24^. Much like Panoptes, ∼50% of PanDA systems were encoded within 10 genes of predicted CBASS systems, frequently as adjacent operons^9^. Also akin to Panoptes, PanDA-adjacent CBASS systems generally encode a 2TM-β or patatin-like phospholipase effector in tandem with Ub-conjugation-like machinery.

### *Ec*PanDA is an antiphage system which defends against phage expressing Acb4

Based on our phylogenetic analyses, we selected a two-gene PanDA operon from *Escherichia coli* MS 185-1 to interrogate (**Figure 1b**). The first gene in this operon, *panS,* encodes a minimal diadenylate cyclase domain that, based on its similarity to characterized DACs, is predicted to synthesize nucleotide signaling molecules^24,25,37^. The second gene, *panE*, encodes a CBASS Cap15/Panoptes OptE-like protein with a SMODS-associating 2TM, β-strand rich (S-2TMβ) domain and two transmembrane domains^37–39^. We expressed the PanDA operon in *E. coli* MG1655 and challenged these bacteria with a diverse set of phages from the BASEL collection^43,44^. The PanDA operon robustly defended against phages from the *Schitoviridae* family. Specifically, PanDA provided over 100-fold protection against Bas69 and over 50-fold protection against the related *E. coli* phage N4 (**Figure 2a**). Interestingly, PanDA did not restrict Bas96, which is also a member of the *Schitoviridae* family. We also found that PanDA restricted phages from the *Straboviridae* family, but to a much lesser degree (**Figure 2a**).

**Fig. 2:**
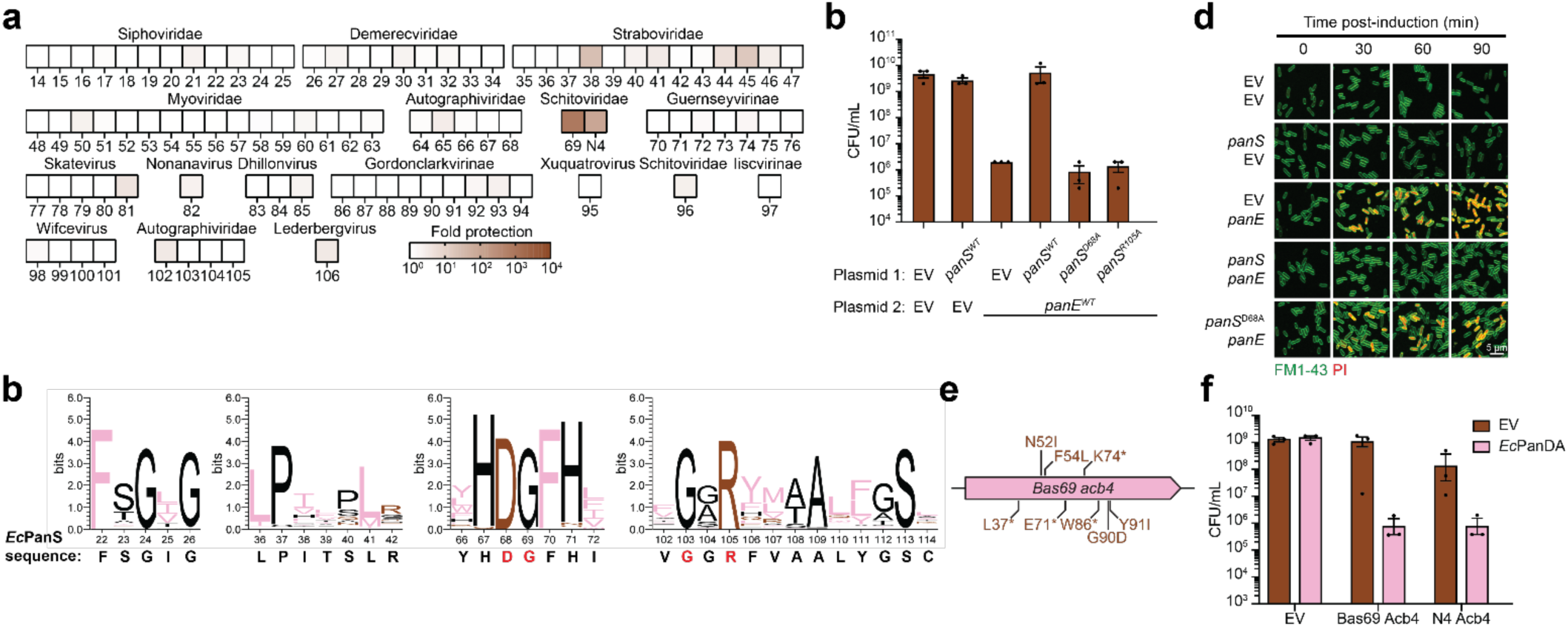
PanDA provides protection from phages expressing the immune evasion gene *acb4*. a) Defense phenotypes of the *Ec*PanDA system expressed in E. coli against a panel of diverse phages. Defense is plotted as heatmap with intense brown representing strong defense. b) Sequence logos generated to illustrate conservation of amino acids across the mDAC clade of diadenylate cyclases. S, G, H, T, A, P are colored black. R, K, D, E, N, Q are colored brown. Y, V, M, C, L, F, I, W are colored pink. *Ec*PanS residues colored in red are 100% conserved across the mDAC clade. c) Dual plasmid expression experiment (no phage) showing toxicity of PanE, necessity of PanS to negate toxicity of PanE, and importance of cyclic nucleotide synthesis (residues) in controlling this. d) Microscopy showcasing propidium iodide staining. e) Depiction of phage Bas69 escaper mutants mapped onto the acb4 gene f) No phage, expression of Bas69 or N4 phage acb4 alone is sufficient to trigger PanDA defense.

We sought to identify residues important for PanS activity and gauge their conservation across PanDA systems and non-immune DACs. We first aligned members of the minimal DAC clade and generated a consensus sequence logo (**Figure 2b**). In terms of *Ec*PanS, D68, G69, G103, R105 were 100% conserved across the alignment. Of note, F22 and F70 were conserved in ∼99.5% of sequences and S113 was conserved in 99.1%. Additionally, an HDGFH motif was conserved in ∼94% of predicted PanS homologs. In an alignment of all curated bacterial DACs on the tree, D68, G69, G103, and R105 again remained fully conserved. We also noted that regions of conservation generally fell on secondary structure elements of the DAC domain (**Extended Data Figure 2e**). We therefore hypothesized that D68A and R105 play key roles in *Ec*PanS activity and targeted D68 and R105 for alanine mutagenesis in cells expressing *Ec*PanDA. Our hypothesis was supported by published structural analyses of *Tma*DisA in which homologous residues (D75 and R108) were found to coordinate metal binding and ribose contacts, respectively^24,45^ (**Figure 2b**).

Given the homology between the Panoptes OptE and PanDA PanE proteins, and that PanS is predicted to synthesize a nucleotide second messenger, we hypothesized that the PanDA system worked through a Panoptes-like mechanism. If this were true, then PanE should inhibit colony formation in the absence of PanS-derived nucleotide products. To test this, we expressed *panS* and *panE* from IPTG- and arabinose-inducible promoters, respectively. *PanS* expression alone had no impact on bacterial growth, but expression of *panE* robustly suppressed colony formation (**Figure 2c**). Inhibition of growth was rescued when *panE* was co-expressed with *panS* in the same strain. Co-expression of *panE* with a *panS* that contained a point mutation in either a conserved metal or ligand binding residue (D68A or R105A), however, could not restore bacterial growth (**Figure 2c**).

We explored the effect of *panE* activation *in vivo* by carrying out the same co-expression experiment but instead monitored the status of the bacteria by laser scanning confocal microscopy. The resulting images, taken after induction with IPTG and arabinose, showed that expression of *panE* alone resulted in propidium iodide (PI) staining starting 30 minutes post-induction, indicative of bacterial membrane permeability (**Figure 2d**). Similar PI staining was observed when *panE* was co-expressed with catalytically dead *panS*^D68A^. Strains that expressed *panS* alone, or both *panS* and *panE*, however, did not show any PI staining, even after an hour post-induction (**Figure 2d**).

To identify the phage component that activates the PanDA system during infection, we generated Bas69 escaper phages on *panDA*-expressing MG1655 *E. coli*. Three candidate escaper phages from each of the six unrelated, but clonal Bas69 lineages were plaque purified and sequenced along with their parent wild-type phages. A majority of the escaper phages encoded mutations in the anti-CBASS 4 gene (*acb4*). The most prevalent type of mutation was single nucleotide polymorphisms (SNP) that resulted in premature stop codons, but we also observed SNPs in *acb4* that led to amino acid substitutions distributed across the coding sequence (**Figure 2e**). Acb4, like the well-characterized anti-CBASS 2 (Acb2), is an anti-defense protein that inhibits CBASS immunity by acting as a nucleotide “sponge,” sequestering cyclic oligonucleotide signaling molecules^46^. Acb4 binds CBASS signaling nucleotides during infection, which blocks downstream effector protein activation. This, in turn, allows the phage to evade CBASS-mediated defense. The finding that escaper phages encoded loss of function mutations in the Acb4 “sponge” anti-defense protein gave credence to our hypothesis that the PanDA system works in a similar way to Panoptes, where anti-defense activity by the phage is a trigger for the system.

We investigated whether expression of Bas69 *acb4* in the absence of any other phage component was sufficient to activate PanDA. As a member of the S-2TMβ family of defense system effector proteins—like OptE of Panoptes—we hypothesized that PanE might also disrupt the bacterial host membrane and restrict colony formation. To test this, we co-expressed the PanDA system with Bas69 *acb4* and measured colony formation. Colony formation was inhibited over 1000-fold when the PanDA operon was expressed only with Bas69 Acb4, but not an empty vector (**Figure 2f**). A similar phenotype was observed when N4 Acb4 was co-expressed with PanDA, restricting colony formation by over 100-fold (**Figure 2f**). These data show that Acb4 is sufficient to activate the PanDA operon. All together, these results support that PanDA defense against phages operates through a Panoptes-like mechanism.

### *Ec*PanS is a minimal DAC which constitutively produces c-di-AMP and cUAMP

To define the structural basis of mDAC activity, we determined the crystal structure of *Ec*PanS (**Figure 3a, Extended Data Table 1**). The 2.14 Å structure (PDB: 37TH) revealed the expected homodimeric complex but unexpectedly arranged around a cyclic-dinucleotide-shaped density (**Figure 3a; Extended Data Figure 2a**). There were four protomers present within the structure (two dimers) which exhibited negligible conformational variation, with superimposition of all protomer pairs yielding an average root mean square deviation (r.m.s.d.) value of ∼0.256 Å. Similar to the DisA protein DAC domain, *Ec*PanS protomers are composed of a seven-stranded β-sheet surrounded by a bundle of four α-helices^24^. Dimers of the DAC domains from *Ec*PanS and *Thermotoga maritima* DisA (*Tma*DisA) aligned well, with superimposition of the two yielding an r.m.s.d. value of 2.429 Å (**Figure 3b**). *Tma*DisA assembles as an octamer by means of two tetramerization events that are dependent on its helical spine domain^24^. Given that *Ec*PanS lacks any additional domains which might facilitate such higher order oligomerization, we suspect that the conserved active site formed between PanS dimers remains the extent of oligomeric state changes in this class of mDACs.

**Fig. 3:**
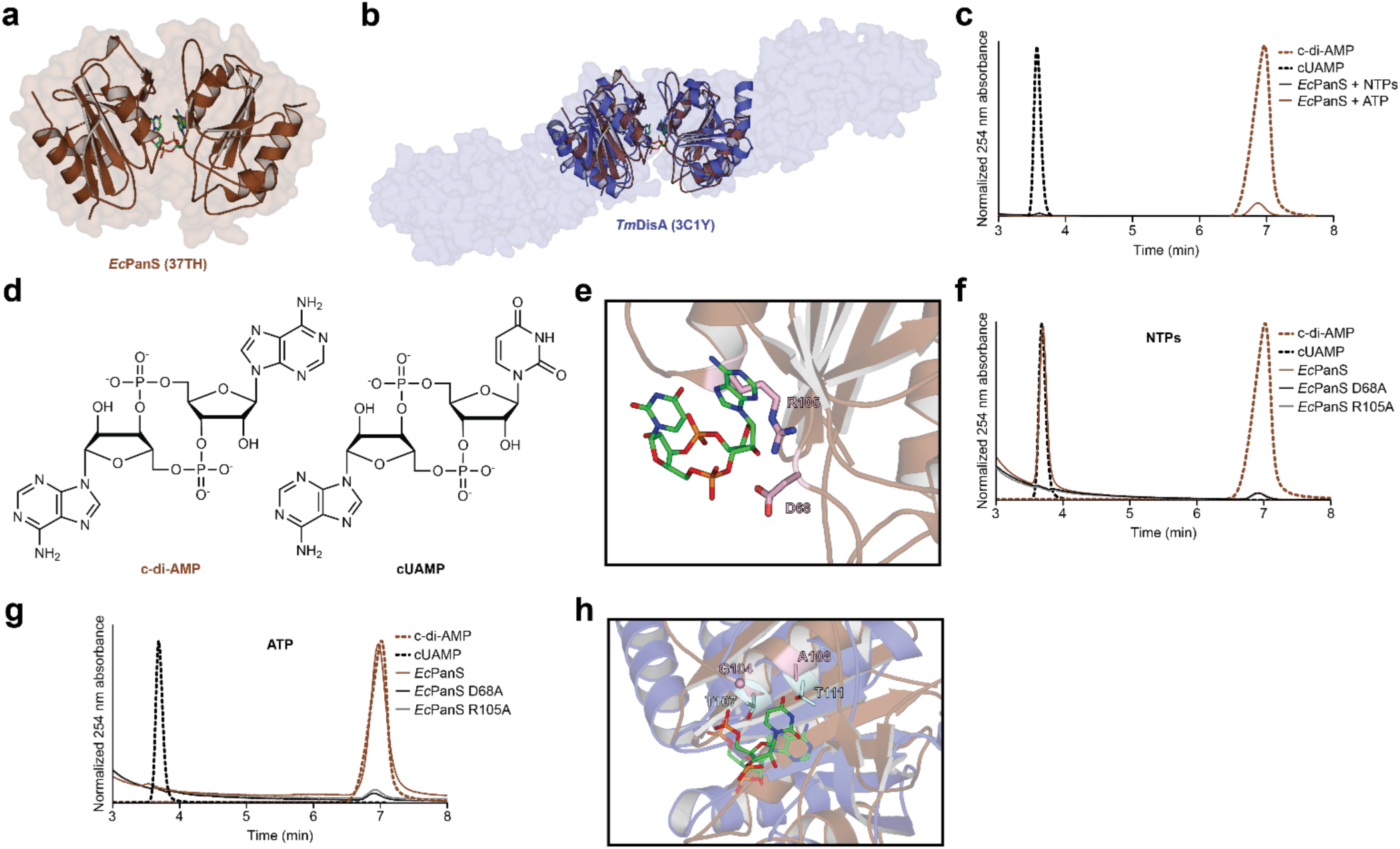
*Ec*PanS constitutively synthesizes c-di-AMP and cUAMP. a) Crystal structure of *Ec*PanS bound to cUAMP (green) at the dimeric interface (37TH). b) Structural superimposition of *Ec*PanS with *Tm*DisA highlights the size difference between most DAC-domain containing proteins and PanDA defense system mDACs. *Tm*DisA in blue with DNA-binding and linker domains shown as a surface model. c) High-performance liquid chromatography data showing *Ec*PanS makes cUAMP in the presence of NTPs and a minor amount of c-di-AMP, or it produces c-di-AMP with ATP alone. Representative of three biological replicates. d) Chemical structures of c-di-AMP and cUAMP. e) Product-bound *Ec*PanS reveals potential residues important for catalysis, specifically D68 and R105 are shown as pink sticks. f, g) High-performance liquid chromatography data traces showing the impact of alanine substitutions on *Ec*PanS activity in the presence of all four NTPs or with ATP, respectively. Representative of three biological replicates. h) Structural comparison of *Tm*DisA (blue) to *Ec*PanS (brown) showing residues responsible for promiscuity of product.

Surprisingly, upon refinement, the dinucleotide was identified as 3′3′-cyclic-UMP-AMP (cUAMP). This was unexpected as no published work has shown DAC production of dinucleotides other than c-di-AMP. Attempts to purify apo *Ec*PanS via anion-exchange chromatography failed as the nucleotide remained present in the structure of the ion-exchanged protein crystalized under different conditions than 37TH (PDB: 37TG, 1.99 Å; **Extended Data Figure 2b**; **Extended Data Table 1**). To better understand *Ec*PanS nucleotide selectivity, we incubated the protein overnight either with an equimolar combination of ribonucleotide triphosphates (ATP, GTP, CTP, UTP) or solely with ATP. Analysis of reaction products via high-performance liquid chromatography (HPLC) showed that *Ec*PanS supplemented with all four NTPs produced a limited quantity of cUAMP and a minute peak for c-di-AMP (**Figure 3c, d**). In contrast, *Ec*PanS supplemented with ATP produced a more substantial c-di-AMP peak, supporting that it is likely the physiologically relevant product. Proteinase K treatment of *Ec*PanS followed by further HPLC analysis confirmed that the protein copurifies with cUAMP, as identified in the crystal structure (**Extended Data Figure 2c).** Production of cUAMP by PanS homologs is not ubiquitous as a similar assay with an *Acinetobacter* PanS (*Ab*PanS) homolog only yielded c-di-AMP even when supplied with all four NTPs (**Extended Data Figure 2d**).

To disentangle the catalytic promiscuity of *Ec*PanS, we incubated alanine mutants of the conserved D68 and R105 residues with either all four NTPs or ATP alone. Further HPLC analysis demonstrated that cyclase activity was inhibited for both mutants with either combination of nucleotides, supporting our hypothesis that these residues are involved in aspects of catalysis separate from nucleotide selectivity (**Figure 3f, g**). A previous structural analysis of *Tma*DisA concluded that the residues T107 and T111 directly interact with the adenine moiety during c-di-AMP synthesis^45^. We therefore aligned *Ec*PanS, *Ab*PanS, *Tma*DisA, and other hits from our tree to identify homologous residues (**Extended Data Figure 2e**). We found that *Ec*PanS possessed G104 and A108 in place of the respective *Tma*DisA residues, while *Ab*PanS encoded A112 and T116 instead. Superimposition of *Ec*PanS and the c-di-AMP*-*bound *Tma*DisA (3C1Y) confirmed that G104 and A108 occupy similar positions within the active site as do T107 and T111 of *Tma*DisA (**Figure 3h**). We therefore suspect that *Ec*PanS synthesis of cUAMP hinges on the greater availability of space within the active site which may accommodate the uracil moiety likely precluded by the respective residues of *Tma*DisA and *Ab*PanS.

### *Ec*PanE is a predicted 2TM-B effector which binds c-di-AMP, but not cUAMP

2TM-β effectors are the most common transmembrane effectors in CBASS systems (denoted as Cap15) and frequently occur in CBASS-like systems such as Panoptes (denoted as OptE)^9,10,47^. Superimposition of crystal structures of the *Yersinia aleksiciae* Cap15 (7N34) and *Vibrio navarensis* OptE (37TF) onto an AlphaFold3 predicted model for the *Ec*PanE β-barrel yield r.m.s.d. values of 0.972Å and 1.817Å, respectively (**Figure 4a**). Like *Ya*Cap15 and *Vn*OptE, *Ec*PanE also possesses a solvent exposed binding pocket which is likely the primary ligand-binding site (**Extended Data Figure 3a**). To compare the binding affinity of *Ec*PanE with the dinucleotide products of *Ec*PanS, we performed isothermal titration calorimetry (ITC) experiments and titrated in either c-di-AMP or cUAMP. A global fit of three replicates for each dinucleotide showed that *Ec*PanE endothermically binds c-di-AMP with a 7.7 ± 0.3 µM affinity while no binding was detected with cUAMP, substantiating that c-di-AMP is likely the primary product of *Ec*PanS (**Figure 4b, c; Extended Data Figure 3b-g).** Immune receptors across diverse taxa generally bind their cognate ligands with an affinity in the nanomolar range, reflective of a very stable binding event capable of effectively propagating downstream immunity^12,48–52^. In contrast, an affinity in the micromolar range is frequently characteristic of binding to non-cognate ligands^12,50–52^. *Ec*PanE binding c-di-AMP with micromolar affinity implies a greater degree of ligand-dissociation which coincides with our hypothesized model that PanDA is a Panoptes-like system which detects dinucleotide sequestration by phage sponge proteins.

**Fig. 4:**
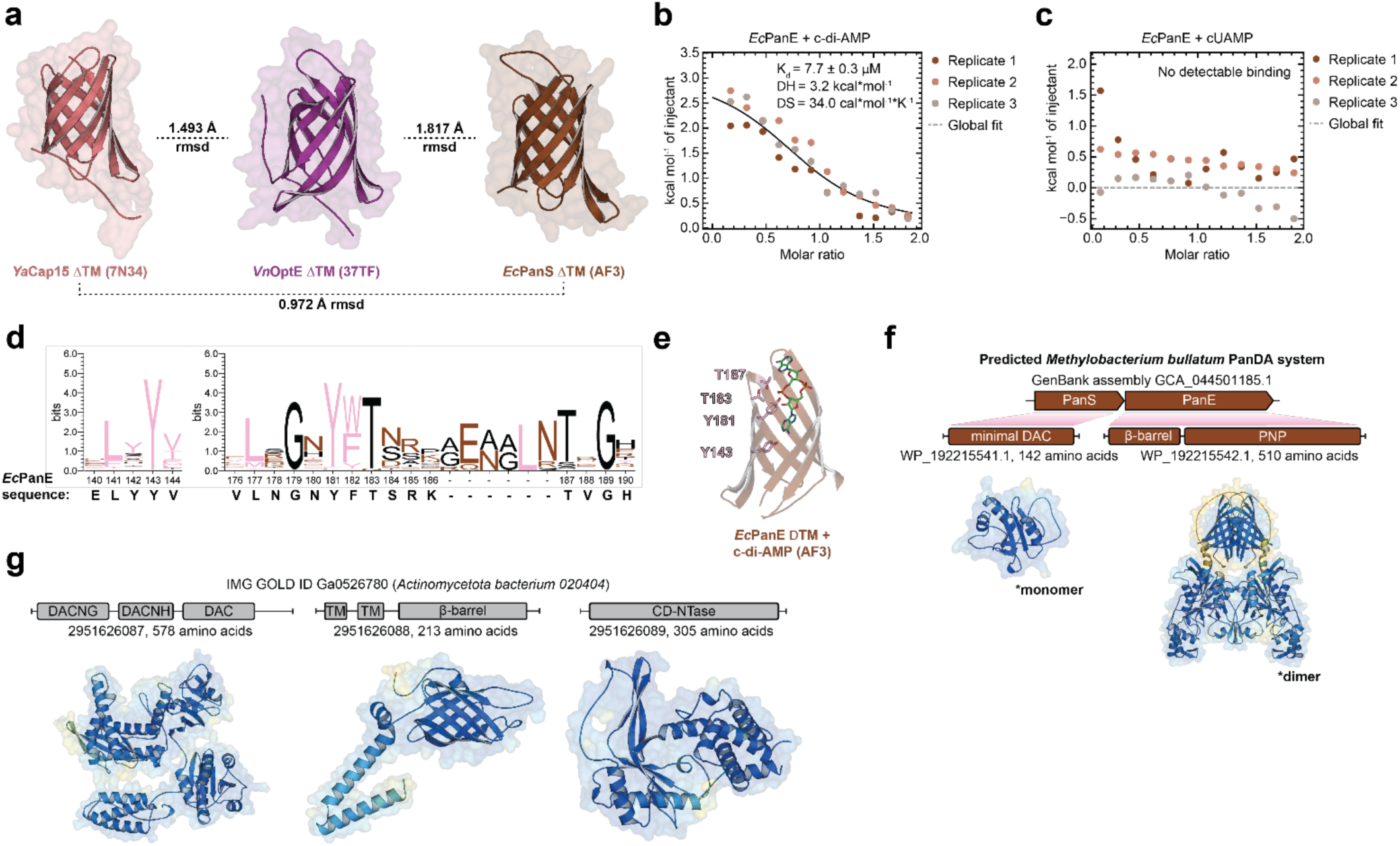
*Ec*PanE binds c-di-AMP through its β-barrel domain. a) Structural comparison of the soluble β-barrel domains of 2TM-β effectors from different bacterial antiphage systems. Left-Published crystal structure of CBASS YaCap15 (PDB: 7N34); Middle-crystal structure of Panoptes *Vn*OptE determined at 2.05 Å (PDB: 37TF, this work); Right-Alphafold3 prediction of *Ec*PanEΔTM. Low rmsd values indicate a high degree of structural similarity amongst these β-barrel proteins. b, c) Isothermal titration calorimetry data for purified *Ec*PanE with -di-AMP or cUAMP, respectively. Each plot shows data from three independent replicates and the associated global fit of the data to extract K_d_ values. d) Sequence logo analysis of conservation of predicted cyclic dinucleotide binding pocket residues in PanE homologs from PanDA defense systems. S, G, H, T, A, P are colored black. R, K, D, E, N, Q are colored brown. Y, V, M, C, L, F, I, W are colored pink. e) Structural visualization of the putative binding pocket and plausible cyclic dinucleotide preference-conferring residues in an *Ec*PanE Alphafold3 model co-folded with c-di-AMP. f) PanDA defense operons contain effectors other than β-barrels with transmembrane domains. Other effectors next to mDACs include a small % of β-barrel domains fused to PNP domains. Alphafold3 predicted structures of a representative operon are shown. g) Other than mDACs having predicted immune function, a separate clade of DACNG-like homologs are also predicted to have immune relevance and are associated with a variety of CBASS systems. A representative DACNG and adjacent CBASS system containing a 2TM-β effector and CD-NTase are shown.

We next sought to identify residues which likely participated in *Ec*PanE binding to c-di-AMP. We gathered the sequences of putative mDAC-associated PanE homologs and aligned them with MAFFT. Sequence analysis revealed several highly conserved residues across the full length of the alignment (**Figure 4d**). To pinpoint residues likely to coordinate adenine selectivity, we co-folded *Ec*PanS with c-di-AMP (via AF3) and examined highly conserved residues (>85% sequence identity) within the binding pocket. Due to their conservation and predicted orientation with respect to the dinucleotide, we noted Y183, Y181, T183, and T187 as the residues most likely to coordinate the adenine moieties of c-di-AMP (**Figure 4e**).

Finally, we also identified other rare DAC-effector pairings during our initial search. In the context of putative PanDA systems, <5% of hits within the mDAC clade were encoded adjacent to effectors composed of a β-barrel appended to a purine nucleoside phosphorylase domain (β-PNP) (**Figure 1a**, **Figure 4f)**. PNP domains are found in effectors of a wide variety of CBASS, pAgo, Avs, Detocs, and other bacterial immune systems where they deplete cellular ATP in response to phage infection^53^. In the context of PanDA, we predict that the PNP domain retains this effector function while the β-barrel again functions as a c-di-AMP receptor. Beyond PanDA systems, we also identified several homologs of the recently categorized CdaG family of DACs encoded adjacent to cGAS/DncV-like nucleotidyltransferases (CD-NTases) and 2TM-β proteins (**Figure 1a**, **Figure 4g**)^26^. CdaG proteins are composed of (from N- to C-termini) DACNG, DACNH, and DAC domains. The function and evolutionary context of DACNG and DACNH domains remain unknown, though the DACNH appears very distantly related to the DAC despite lacking any of the conserved catalytic residues^26^. The adjacency of two protein folds which each synthesize a cyclic dinucleotide second messenger may function to widen the sensitivity of an antiphage system, though it remains to be seen if the shared effector can respond to the products of both cyclases.

### Acb4 activates PanDA by sequestering c-di-AMP

SP01 phage encode an Acb4 homolog which was recently shown to bind and sequester CBASS-derived cyclic dinucleotides, namely 3′3′-c-GMP-AMP, with affinities in the nanomolar range^54^. Other phage sponges which perturb cyclic nucleotide pools utilized in bacterial immunity also bind their cognate ligands with affinities in the nanomolar range^55–58^. To verify that Acb4 homologs belonging to PanDA-sensitive phage are similarly able to bind c-di-AMP, we performed ITC on the phage N4 Acb4 homolog while titrating in either of the dinucleotides produced by *Ec*PanS. We measured that N4 Acb4 binds c-di-AMP with an affinity of 111 ± 13 nM (**Figure 5a**; **Extended Data Figure 3h, i**). In contrast, the sponge bound cUAMP with an affinity of 13.7 ± 1.4 µM, roughly two orders of magnitude weaker than c-di-AMP (**Figure 5b**; **Extended Data Figure 3j, k**). We confirmed that these ITC-measured affinities are reflected by Acb4 binding of *Ec*PanS products using an HPLC assay in which sponge was added to reactions containing the cyclase incubated with either NTPs or ATP alone. After incubation with N4 Acb4, protein within the reactions was digested with Proteinase K (ProK) to confirm the release of the suspected dinucleotide. The addition of Acb4 to reactions containing all four NTPs incompletely sequestered the cUAMP produced by *Ec*PanS while the sponge was able to completely deplete c-di-AMP generated by *Ec*PanS incubated with ATP alone (**Figure 5c, d**). In both sample conditions, digestion by ProK released the majority of the respective dinucleotide.

**Fig. 5:**
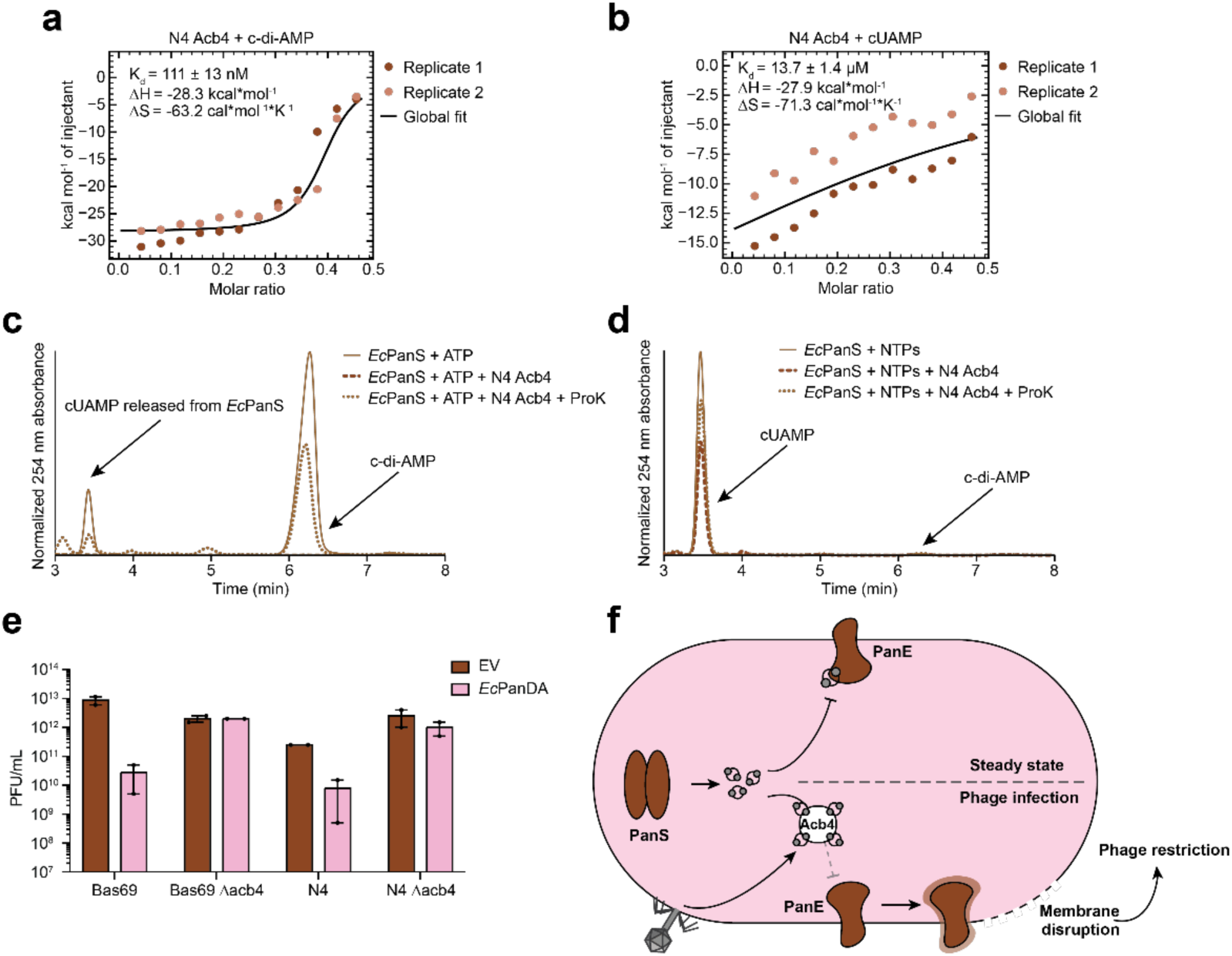
Phage N4 Acb4 sequesters c-di-AMP and is sufficient to trigger PanDA defense. a, b) Isothermal titration calorimetry data for purified phage N4 Acb4 incubated with c-di-AMP and cUAMP, respectively. Each plot shows data from two independent replicates and associated global fit of the data to extract K_d_ values. c, d) High-performance liquid chromatography data for enzymatic reactions of *Ec*PanS with NTPs or with ATP and treated with phage N4 Acb4 and Proteinase-K. Representative of three biological replicates. e) Plaque forming units as measured for wildtype and Acb4 knockout phages infecting *E. coli* expressing *Ec*PanDA (pink) or empty vector control (brown). Two biological replicates, error expressed as SEM. f) Generalized model of PanDA defense in bacteria.

Finally, we substantiated that Acb4 specifically was necessary for PanDA-mediated defense by infecting empty vector or PanDA-expressing bacteria with either wild-type or Δ*acb4* Bas69 and N4 phage. The wild-type phage were restricted by PanDA but replicated well on an empty vector-expressing strain (**Figure 5e**). Bas69 and N4 phage lacking *acb4*, however, were able to replicate efficiently on both empty vector- and PanDA-expressing strains (**Figure 5e**). This data again suggest that the *acb4* gene is necessary for PanDA-mediated defense. In turn, we assert that the mechanism by which PanDA defends against Acb4-encoding phage entails a Panoptes-like detection of c-di-AMP sequestering by phage sponge proteins, leading to membrane disruption and phage restriction (**Figure 5f**).

## DISCUSSION

In this study we discovered a novel antiphage system which we named PanDA, incorporating a minimal DAC domain (PanS) and a toxic 2TM-β effector (PanE). DAC domain-containing proteins have long been characterized by their appended regulatory domains which function to modulate essential cyclase activity in the context of varied nonimmune processes such as responses to pH, osmotic, and temperature stresses^26^. In PanDA, PanS was found to lack a regulatory domain which instead implies constitutive activity. Indeed, we found that PanS constitutively synthesizes c-di-AMP to repress PanE, highlighting the mechanistic similarity to systems such as Panoptes and Hailong^9,10,59,60^. PanDA defends against phage within the *Schitoviridae* family, specifically those encoding homologs of the sponge protein Acb4, but does not defend against those lacking an endogenous Acb4 such as Bas96. Acb4 binds c-di-AMP with nanomolar affinity, similar to other sponges used by phage to evade bacterial immunity^9,10,55–58^. PanDA additionally provided lesser defense against *Straboviridae* phage, likely due to their encoding homologs of Acb2, a similar phage sponge which was recently shown to activate the Panoptes system^9,10^. We establish that sequestration of c-di-AMP by Acb4 releases its repression of PanE, leading to membrane permeabilization and restriction of phage replication.

Previous work has focused on a small subset of the extremely diverse families of bacterial DAC domain-containing proteins. In our initial search, we charted the phylogenetic relatedness of both previously characterized and novel DAC domain organizations. In the process, we identified a large sequence divergent clade of PanS homologs, a smaller subset of likely immune proteins belonging to the CdaG family, as well as several non-immune DACs with relatives reported on in the literature^26^. Our search revealed that non-immune DACs are very widely distributed across bacterial phyla, bringing into question whether a system such as PanDA could coexist in the same genome as a non-immune DAC without destabilizing cellular homeostasis. Indeed, we found no instance of a PanS encoded within the same genome as a non-immune DAC, though a handful were found in the same genome as the predicted CdaG proteins also represented on our tree. This supported the notion of a mutual exclusivity of DAC activity in and outside of immunity within the context of a single organism. It remains an open question whether phage encoding proteins which perturb cellular c-di-AMP pools infect bacteria which incorporate DACs in non-immune processes. This may represent a yet unnoticed means for phage to destabilize a bacterial host during infection.

PanDA belongs to a subset of systems which exapted widespread non-immune protein folds for antiphage defense. It can be reasoned that phage may, on a viral population-level scale, face less selective pressure to evolve and acquire proteins which antagonize the products of bacterial signaling enzymes which are generally restricted to non-immune processes. As such, the acquisition of such non-immune protein domains for use in bacterial immunity may embody a bacterial ace in the hole against phage infection. Future work may demonstrate that other protein domains, such as the GGDEFs, have been similarly exapted for use in antiphage systems in response to comparably limited viral targeting of core host processes.

The substantial diversity of DAC regulatory domains, due in some part to the speed of bacterial evolution and the tendency of horizontal gene transfer between species, obfuscates the evolutionary history of the cyclase domain. From our tree we can reason that PanS is more closely related to membrane-bound DACs found in the adjacent clade, perhaps indicating that the minimal DAC was formed upon the loss of its transmembrane domain. In contrast, the likely immune CdaG homologs we identified are nested within a clade containing several proteins with no homology to characterized DACs in the literature. The functional role of the DACNG domain, DACNH domain, and C-terminal helix bundle present within CdaG proteins would shed a light on evolutionary origin of members of this clade and represents an exciting new frontier in the exaptation of the DAC domain for use in bacterial immunity.

## METHODS

### Bioinformatics analysis and phylogenetic tree generation

Separate searches were conducted with MMSeqs2 and PSI-BLASTp to identify the DACs examined in this study^61,62^. Searches were seeded with DACs representative of the majority of families recently described by Michael Y. Galperin in the Journal of Bacteriology (2023)^26^. Searches were conducted of a database assembled from the proteomes of approximately 300,000 publicly available bacterial genomes retrieved from the NCBI and IMG databases. 10 iterations of both searches were performed with parameters to ensure maximum sensitivity, query coverage set to .5, and e= 1e-5. A non-sequence-redundant list of hits was generated by retaining the highest scoring hit from each set of hits with identical sequences. An outgroup composed of 50 diguanylate cyclases was added and sequences were aligned via MAFFT using the FFT-NS-I method^63^. A maximum likelihood tree was built with IQTree2 including parameters for ModelFinder Plus, SH-aLRT branch testing with 1000 replicates, and UFBoot2 ultrafast bootstrapping with 1000 replicates^64^. Sequences less than 100 amino acids in length were trimmed to produce the final tree.

Protein neighborhood analysis was conducted by first collecting the 10 proteins proceeding and following each of the DACs identified in the above search. Each neighbor was searched with hmmscan against the PfamA database and with hhblits against the PDB70 database^65,66^.

### Bacterial strains and growth conditions

All bacterial cultures were grown in 3.5 mL of media in 14 mL culture tubes shaking at 220 rpm at 37 °C, unless otherwise indicated. “Overnight” cultures are those that were grown for 16–20 hours following inoculation from a single colony or glycerol stock. Where applicable, culture media was supplemented with chloramphenicol (20 µg/mL) and/or carbenicillin (100 µg/mL) for plasmid maintenance or strain selection. *E. coli* OmniPir^67^ was used for cloning and storage of plasmids and *E. coli* MG1655 (CGSC6300) was used for all microscopy and phage and colony formation experiments.

All *E. coli* cultures used for cloning, strain construction, perform phage amplification, phage infection assays, and colony formation assays were grown in LB medium (1% tryptone, 0.5% yeast extract, and 0.5% NaCl). All strains were frozen for long-term storage in LB plus 30% glycerol (v/v) at −70 °C. Strains used for microscopy were cultivated in “MMCG” minimal medium (47.8 mM Na_2_HPO_4_, 22 mM KH_2_PO_4_, 18.7 mM NH_4_Cl, 8.6 mM NaCl, 22.2 mM glucose, 2 mM MgSO_4_, 100 mM CaCl_2_, 3 mM thiamine, Trace Metals at 0.1× (Trace Metals mixture T1001, Teknova, final concentration: 8.3 μM FeCl_3_, 2.7 μM CaCl_2_, 1.4 μM MnCl_2_, 1.8 μM ZnSO_4_, 370 nM CoCl_2_, 250 nM CuCl_2_, 350 nM NiCl_2_, 240 nM Na_2_MoO_4_, 200 nM Na_2_SeO_4_, 200 nM H_3_BO_3_)). When a strain with two plasmids was cultivated in MMCG medium, bacteria were grown in carbenicillin (20 μg/mL) and chloramphenicol (4 μg/mL). When growing strains that required induction, 500 μM IPTG or 0.2% arabinose was used to induce, as appropriate. LB and MMCG agar plates contain 1.6% agar and media components described above.

### Molecular cloning and plasmid construction

Cloning and plasmid construction were performed as previously described^68^. Briefly, genes of interest were amplified from previously constructed plasmids or phage genomic DNA using Q5 Hot Start High Fidelity Master Mix (NEB, M0494L). Gene inserts were flanked by at least 18 homologous base pairs to the vector backbone outside of the restriction digest sites. Ligation of genes into the digested, linearized backbone vector was done using modified Gibson Assembly^69^ with HiFi DNA Assembly Master Mix (NEB, E2621L). Gibson assemblies were transformed by electroporation into competent OmniPir and plated onto LB (1.6 % agar) plates with appropriate antibiotics to select for successfully transformed bacteria. Point mutations in *PanS* were generated by amplifying the gene of interest in two parts from a plasmid template, with the desired mutation occurring in the overlapping region between the two amplicons.

For the *PanSE* operon in a pLOCO3 backbone, complete vectors with the indicated operons were generated as ValueGENEs (Genewiz). The pLOCO3 vector (including sfGFP) was initially constructed using Gibson assembly to join and circularize two FragmentGENEs, one with the pLOCO3 backbone, and one with the sfGFP gene.

For all vectors using the pTACxc backbone, pAW1608 was amplified and purified from OmniPir. Purified plasmid was then linearized using BmtI-HF and NotI-HF, or BmtI-HF and PstI-HF. Gibson ligation was used to circularize the plasmid with the new insert.

For all vectors using the pBAD30 backbone, pAW1640 was amplified and purified from OmniPir. Purified plasmid was then linearized using EcoRI-HF and NotI-HF. Gibson ligation was used to circularize the plasmid with the new insert.

Synthetic dsDNA fragments for each gene of interest for protein expression and purification were produced in codon optimized form (IDT) and cloned into a pET16-N6xHis-SUMO2 custom vector using Gibson assembly. BamHI and NotI restriction enzymes were used to linearize the plasmid prior to assembly. Around-the-horn mutagenic primers were used to generate PanS mutants. All plasmid sequences were verified using Sanger or Oxford Nanopore Sequencing (Genewiz or Plasmidsaurus).

### Bacteriophage amplification and storage

Information on the complete BASEL collection used in this study is reported in Maffei et al. 2021^43^ and Humolli et al. 2025^44^. Phage lysates were generated via plate amplification using a modified double agar overlay^70^. Briefly, 400 µL of mid-log MG1655 were mixed with 3.5 mL LB soft agar mix (LB media with 0.35 % agar and 10 mM MgCl_2_, 10mM CaCl_2_, and 0.1 mM MnCl_2_) and 100-1,000 PFU and poured on top of a LB (1.6% agar) plate. Plates were then incubated overnight at 37 °C. Phages were collected by adding 5 mL of SM buffer (100 mM NaCl, 8 mM MgSO_4_, 50 mM Tris-HCl pH 7.5, 0.01% gelatin) to the plate and incubating for 1 hour at room temperature. To increase phage titers, the top agar overlay was scraped and harvested along with the SM buffer. The SM buffer and top agar mixture was centrifuged at 4,000 × g for 10 minutes and the supernatant was transferred to a new tube. The resulting liquid was treated with 10 drops of chloroform, followed by vortexing, to remove any remaining viable bacteria. Amplified phages were stored at 4 °C in SM buffer.

### Bacteriophage infection assays

Phage titer quantifications and phage infection assays were performed using a modified double agar overlay technique^70^. Strains containing the indicated plasmids were cultivated overnight in LB medium (including appropriate antibiotics) and were diluted 1:10 or 1:100 in fresh medium the following day. The bacteria were grown until they reached mid-logarithmic phase (OD_600_ 0.1-0.8). 400 µL of mid-log bacteria were mixed with 3.5 ml 0.35 % LB agar (plus 10 mM MgCl_2_, 10mM CaCl_2_, and 0.1 mM MnCl_2_) and poured on top of a LB (1.6% agar) plate, respectively. The top agar was allowed to cool and solidify for 12 minutes. Once cooled, 2 μL of a phage 10-fold serial dilution series was spotted onto the soft agar overlay and allowed to dry, after which the plates were incubated at 37 °C overnight. Plates were imaged ∼16-20 hr after infection.

The resulting phage titer was quantified in PFU/mL for each phage lysate tested. PFU were counted based on the lowest phage dilution spot with individual, quantifiable PFU. The dilution at that spot was used to calculate the PFU/mL. When there was a hazy zone of clearance rather than identifiable plaques, the lowest phage concentration at which this was seen was counted as ten plaques. When there was no clearance observed, the least dilute spot was counted as 0.9 plaques, and this was used as the limit of detection for the assay.

### Bacteriophage escaper generation and analysis

Bas69 escaper phages were generated from six unrelated, clonal Bas69 lysates (“parents”) that were separately plate amplified on wild-type *E. coli* MG1655. To make the escaper phages (“daughters”), 400 µL of mid-log bacteria expressing the *PanSE* operon (in LB plus 100 µg/mL carbenicillin) was mixed with 100 µL of parent Bas69 lysate and 3.5 mL LB top agar and poured onto a LB agar plate. The plate was allowed to dry and was incubated overnight at 37 °C. The next day, three single escaper plaques were individually isolated from each parent Bas69 plate using a Pasteur pipette, soaked in 500 µL SM buffer in an Eppendorf tube, and sterilized with 3-5 drops of chloroform, followed by vortexing. A dilution series of each Bas69 escaper phage was spot plated onto *E. coli* MG1655 expressing the PanDA system to confirm replication in the presence of *PanSE*. From this plate, single plaques from each escaper were individually purified and plate amplified (as described in the phage amplification and storage protocol above) before storage.

The genomes of the parent and escaper Bas69 phages were purified as previously described^71^. To do this, 450 mL of phage lysate (>10^8^ PFU/mL) was mixed with 50 µL 10× DNAse I buffer (100 mM Tris-HCl pH 7.6, 25 mM MgCl_2_, and 5 mM CaCl_2_) and treated with DNAse I (final concentration 2 × 10^-3^ U/µL) and RNAse A (final concentration 2 × 10^-2^ mg/mL). This mixture was incubated for 1.5 hours at 37 °C to remove extracellular nucleic acids. After, EDTA was added to a final concentration 20 mM to stop the reaction. Each parent and escaper phage genome was then isolated and purified using the Qiagen DNeasy cleanup kit, starting at the proteinase K digestion step^71^.

Sequencing libraries of the purified phage escaper genomes were generated using the tagmentation-and PCR-based Illumina DNA Prep kit and custom IDT 10bp unique dual indices (UDI) with a target insert size of 280 bp. Sequencing was performed on an Illumina NovaSeq X Plus sequencer in a multiplexed shared-flow-cell runs, producing 2 x 151bp paired-end reads (SeqCenter, LLC). Bcl-convert (Illumina, v4.2.4) was used for demultiplexing, quality control, and adapter trimming. The resulting reads were mapped to Genome accession MZ501049.1 using Geneious software’s Map to Reference feature. The Geneious feature “Find Variations/SNPs” was used to identify variants in daughter phage genomes. Called variants were identified as escaper mutations if they were present in ≥75% of reads and were not present in parent phage genomes.

Targeted sequencing of the *acb4* gene locus in the parent and daughter Bas69 phages was done using PCR amplification (forward primer oAES0163: ACATCCCAACCACGAAGTTTG; reverse primer oAES0164: GGATGTGTAGCGAGTGAAG) followed by Sanger sequencing (Azenta). The resulting reads were mapped to Genome accession MZ501049.1 using Geneious software’s Map to Reference feature. The Geneious feature “Find Variations/SNPs” was used to identify variants in daughter phage genomes. These called variants were identified as escaper mutations if they were not present in parent phage genomes.

### Colony formation assays for bacterial growth inhibition analysis

Bacterial growth inhibition was tested using colony formation assays. Bacterial strains with indicated plasmids were grown overnight in LB media plus appropriate antibiotics. The cultures were then 10-fold serially diluted in fresh LB media (without antibiotics) and 5 µL of each dilution was spotted onto an LB agar plate containing the appropriate antibiotics, as well as IPTG (500 µM; induced condition) as indicated. Data were collected using LB media with appropriate antibiotics, with or without glucose (0.2% w/v; uninduced condition), IPTG (500 µM; induced condition), or arabinose (0.2% w/v; induced condition), as indicated. After the spotted bacteria was allowed to dry, plates were incubated at 37 °C for ∼16–18 hr. Growth inhibition was quantified the next day by counting the number of colony forming units (CFU) of the lowest dilution that had individual colonies. When no individual colonies could be counted, the lowest bacterial concentration at which growth was observed was counted as ten CFU. In instances where no growth was visible, the least dilute spot was counted as 0.9 CFU and used as the limit of detection.

### Laser scanning confocal microscopy

Strains were grown overnight in MMCG plus appropriate antibiotics. The next day, cells were diluted 1:10 in fresh MMCG media. Strains were grown to mid-logarithmic phase (OD_600_ ∼0.7) and induced with 500 µM IPTG and 0.2% arabinose. Following 30 minutes of incubation the strains were OD normalized to 0.7 in a volume of 100 μl with fresh MMCG medium and stained with 25 µg ml^−1^ of DAPI, 5 µg ml^−1^ of FM1-43, and 5 µM of propidium iodide by incubating samples with dyes for 2 min at room temperature. A total of 5 µl of each sample was pipetted onto an MMCG imaging pad containing 500 µM of IPTG and 0.2% arabinose and allowed to dry for about 5 min before imaging. Samples were imaged using a Nikon AXR confocal microscope system. Images were acquired using a 100x oil objective and 1.45 numerical aperture. Images have a pixel size of 0.09 µm/pixel with a final image size of 512 x 512 pixels. The images were acquired where DAPI signal was measured using an excitation of 405 nm and an emission of 418-475 nm, FM1-43 signal was measured using an excitation of 488 nm and an emission of 500-550 nm, and PI signal was measured using an excitation of 561 and an emission of 575-650 nm. All images were acquired using the same laser power. For each strain condition and time point, a 3 × 3 image scan was collected; representative areas in these scans are shown in **Figure 2d**.

### Recombinant protein expression and purification

Plasmids were transformed into competent BL21-CodonPlus(DE3)-RIL *E. coli* cells using heat shock, then plated on MDG media (1.5 % agar, 0.5% glucose, 25 mM Na_2_HPO_4_, 25 mM KH_2_PO_4_, 50 mM NH_4_Cl, 5 mM Na_2_SO_4_, 2 mM MgSO_4_, 0.25% aspartic acid, 100 μg/mL ampicillin, 34 μg/mL chloramphenicol, and trace metals mix [25 μM FeCl_3_×6H_2_O, 10 μM CaCl_2_×2H_2_O, 5 μM MnCl_2_×4H_2_O, 5 μM ZnSO_4_×7H_2_O, 1 μM CoCl_2_×6H_2_O, 1 μM CuCl_2_×2H_2_O, 1 μM NiCl_2_×6H_2_O, 1 μM Na_2_MoO_4_×2H_2_O, 1 μM Na_2_SeO_3_, 1 μM H_3_BO_3_]) and incubated overnight at 37 °C. The following day, 3 colonies were picked and inoculated into 30 mL of liquid MDG liquid media with 100 μg/mL ampicillin and 34 μg/mL chloramphenicol. These small cultures were shaken overnight at 230 RPM at 37 °C, then inoculated into 1 L of M9ZB media (47.8 mM Na_2_HPO_4_, 22 mM KH_2_PO_4_, 18.7 mM NH_4_Cl, 85.6 mM NaCl, 1% Casamino acids (VWR), 0.5% v/v glycerol, 2 mM MgSO_4_, trace metals, 100 μg/mL ampicillin, and 34 μg/mL chloramphenicol) and shaken at 230 RPM and 37 °C until reaching an OD_600_ greater than 2.5. Cultures were then incubated with ITPG to 500 µM overnight at 230 RPM and 16 °C to promote protein expression. After overnight induction, cultures were briefly centrifuged and the resultant pellets were resuspended in 30 mL of 4 °C PBS using an orbital shaker for 20 minutes, then centrifuged again. PBS was decanted, and samples were flash frozen with liquid N_2_ and stored at -80 °C until purification.

Pellets were resuspended in 60 mL of lysis buffer (20 mM HEPES pH 7.5, 400 mM NaCl, 10% v/v glycerol, 30 mM imidazole, and 1 mM dithiothreitol (DTT)), then lysed by sonication at 70% amplitude, with 10 seconds on followed by 20 seconds off, resulting in a total of 2.5 minutes on time. This sonication procedure was performed twice. The resulting samples were centrifuged for 30 minutes at 14,000 RPM and 4 °C in the JA-17 rotor to isolate sonicated lysates from cellular debris. Protein was then purified using immobilized metal affinity chromatography. Clarified lysate was decanted and applied to 4 mL of packed Ni-NTA resin. The resin was then washed sequentially with 20 mL of lysis buffer, 70 mL of wash buffer (lysis buffer supplemented to 1.0 M NaCl), and again with 35 mL of lysis buffer. The protein was eluted in 20 mL of elution buffer (lysis buffer supplemented to 300 mM imidazole) and then transferred to 10 kDa molecular weight cut-off dialysis tubing and dialyzed overnight at 4 °C into dialysis buffer (20 mM HEPES pH 7.5, 250 mM KCl, 1 mM DTT, 5% glycerol). 6×His-hSENP2 (D364–L589, M497A) was added to the protein samples (produced in-house; final concentration of 1:100 hSENP2:protein w/w) to cleave the 6×His-SUMO-tag during the dialysis step. The following day, the dialyzed eluent was concentrated to less than 1 mL using 10 kDa cut-off Amicon filter-concentrators (Sigma). Concentrated samples were then further purified through size-exclusion chromatography using a Hi-Load 16/600 Superdex 75pg column (Cytiva) equilibrated with 20 mM HEPES pH 7.5, 250 mM KCl, and 1 mM TCEP. Fractions containing the protein of interest were verified through SDS-PAGE, then further concentrated to greater than 10 mg/mL. Protein concentrations were measured spectrophotometrically before being flash frozen in liquid N_2_ and stored at -80 °C until needed.

### Protein crystallization and structure determination

*Ec*PanS and *Vn*OptEΔTM crystals were obtained at room temperature using the hanging-drop vapor diffusion method. Briefly, following initial sparse screen crystallization attempts (96-well plates, nanovolume droplets set by Mosquito robot) larger drops containing a 1-to-1 mixture of purified protein (either with or without ion exchange chromatography) and reservoir solution were prepared manually in EasyXtal 15-well plates (NeXtal). Crystal growth was generally observed after 24-72 hours. Crystals for apo *Vn*OptEΔTM grew in the presence of .225M MgCl_2_, .1M Tris pH 8.5, and 18% PEG4000 at 10 mg/mL and were cryoprotected with addition of 10% ethylene glycol. Crystals for *Ec*PanS (no IEX) grew in the presence of 4M sodium formate at 7 mg/mL and were not cryoprotected. Crystals for *Ec*PanS (with IEX) grew in the presence of .1M HEPES sodium salt pH 7.5, 1.5M lithium sulfate at 7 mg/mL and were cryoprotected with addition of 25% glycerol. Cryoprotected crystals were flash frozen in liquid N_2_ and shipped for remote data collection at the Advanced Light Source (ALS, Berkeley, USA).

X-ray diffraction data were collected at ALS Beams 5.0.1, 5.0.2, and 5.0.3. for *Ec*PanS (with IEX), *Ec*PanS (no IEX) and *Vn*OptEΔTM, respectively. Beamline 5.0.1 data were collected at 100 K using a Pilatus3 2M 25 Hz detector at a fixed energy of 12.7 keV. Beamline 5.0.2 data were collected at 100 K using a Pilatus3 6M 25 Hz detector at an energy of ∼12.7 keV. Beamline 5.0.3 data were collected at 100 K using a Pilatus3 2M detector at a fixed energy of 12.7 keV. Data were automatically processed using XDS (X-ray Detector Software) including POINTLESS and AIMLESS for scaling and space group assignment. Molecular replacement was conducted using the Phaser-MR program within PHENIX (version 2.1-6048) with template inputs composed of predicted structural models generated using Alphafold3. Structures were manually repositioned in Coot (version 1.3.1) and iteratively refined with phenix.refine.

For both *Ec*PanS structures, the Elbow tool in PHENIX was used to generate ligand restraints for 3ʹ,3ʹ-cUAMP and the ligand was manually placed into clear electron density and included in refinement (SMILES code: Nc1ncnc2c1ncn2C1OC2COP(=O)(O)OC3C(COP(=O)(O)OC2C1O)OC(C3O)N1C=CC(=O)NC1=O). Final refined structures had the following Ramachandran statistics-*Vn*OptE: 97.44 % favored, 2.56 % allowed, 0.00 % outlier; *Ec*PanS (with IEX): 98.04 % favored, 1.78 % allowed, 0.00 % outlier; *Ec*PanS (no IEX): 97.89 % favored, 2.11 % allowed, 0.00 % outlier. A polder map omit map was generated for cUAMP in Phenix and is presented at 3 σ (**Extended Data Figure 2a**). Full data collection and refinement statistics are summarized in **Extended Data Table 1**. Protein models were visualized and analyzed using PyMOL (version 3.1.8). Atomic coordinates for *Ec*PanS (no IEX), *Ec*PanS (with IEX), and *Vn*OptEΔTM are deposited at the PDB under IDs 37TH, 37TG, and 37TF, respectively.

### High-performance liquid chromatography

Enzymatic reactions were prepared in 150 µL volumes with 500 µM of the respective ribonucleotide triphosphates (ATP, GTP, CTP, UTP), 1 mM MnCl_2_, 100 mM KCl, 20 mM HEPES-KOH (pH 7.5), 20 mM CAPSO (pH 9.4), and 1 µM protein. Reactions were incubated at 37°C overnight (16-20 h) then transferred to 10 kDa cut-off filters and centrifuged for 20 minutes at 21,130 rcf. 10 µL sample volumes were injected onto an Agilent 1200 Series HPLC equipped with a 4.6 x 150 mm and 5 µM particle-size Zorbax Bonus-RP column using an isocratic elution method. Samples were run in a 97% 50 mM NaH_2_PO_4_ (pH 6.8) and 3% acetonitrile buffer system held at 40°C. Reaction component separation was monitored at 254 nm using an UV absorbance detector.

Reactions gauging Acb4 sequestering of PanS were prepared in 100 µL volumes with 500 µM of the respective ribonucleotide triphosphates (ATP, GTP, CTP, UTP), 1 mM MnCl_2_, 100 mM KCl, 20 mM HEPES-KOH (pH 7.5), 20 mM CAPSO (pH 9.4), and 100 µM of wild-type PanS. Reactions were incubated at 37°C overnight (16-20 hrs), after which 100 µM Acb4 was added. Samples were incubated for an additional 1 h at 37°C. When appropriate, reactions were supplemented with 1 µL of 20 mg ml^-1^ proteinase K (NEB, catalogue no. P8107S) and all reactions were further incubated for 1 hr at 37°C. Samples were transferred to 10 kDa cut-off filters and centrifuged for 20 min at 21,130 rcf. Filtered reactions were then analyzed via HPLC as described above.

### Isothermal titration calorimetry

For ITC data collection we used a Malvern Microcal PEAQ-ITC instrument. The experimental method was a single 0.4 μl injection, followed by 12 injections of 3 μl each. The initial injection had a 0.8 second duration and the remaining 12 had a 6 second injection durations. Spacing between all injections was 150 seconds. Reference power was 41.9 μW, stir speed was 750 rpm, and the initial delay was set to 60 seconds. For *Ec*PanE, 400 µM cyclic dinucleotide was titrated into 40 µM protein. For N4 Acb4, 100 µM cyclic dinucleotide was titrated into 40 µM protein. Preliminary analysis was conducted within the MicroCal PEAQ-ITC Analysis Software platform followed by data normalization and figure generation using *pytc*^72^.

### Synthetic nucleotide ligands

Synthetic cyclic dinucleotide ligands used for HPLC and ITC experiments were purchased from Biolog Life Science Institute: 3′3′-c-di-AMP (catalogue no. C 088); 3′3′-cUAMP (catalogue no. C 357).

### Accession numbers

The crystal structure data for *Ec*PanS, *Ec*PanS (with IEX), and *Vn*OptSΔTM have been deposited in the PDB (37TH, 37TG, 37TF, respectively).

## ACKNOWLEDGEMENTS

We thank the members of the Morehouse and Whiteley labs for their critical feedback and advice when developing this work. We thank Dr. Celia Goulding for the shared use of ITC and Mosquito instrumentation. We additionally thank S.N., R.Y., W.P., S.B., E.D., K.S., and M.M.. We thank the Shared Instruments Pool (RRID: SCR_018986) of the Department of Biochemistry at the University of Colorado Boulder for providing access to the Avanti JXN-26 Super Speed centrifuges and rotors, which are funded by National Institutes of Health (NIH) Grant R24OD033699-01. Use of the Nikon A1R microscope in the BioFrontiers Institute’s Advanced Light Microscopy Core (RRID: SCR_018302) was supported by NIST-CU Cooperative Agreement award number 70NANB15H226. Beamlines 5.0.1, 5.0.2, and 5.0.3 of the Advanced Light Source, a DOE Office of Science User Facility under Contract No. DE-AC02-05CH11231, are supported in part by the ALS-ENABLE program funded by the National Institutes of Health, National Institute of General Medical Sciences, grant P30 GM124169-01. The Pilatus detector on beamline 5.0.1 was funded under NIH grant S10OD026941. We thank the staff and affiliate scientists who assist with remote data collection procedures at these beams.

