## Extended Data Figures 1-3 for "Minimal diadenylate cyclases have been co-opted to detect phage immune evasion"

*= equal contribution †= co-corresponding

**EXTENDED DATA FIGURES 1-3**

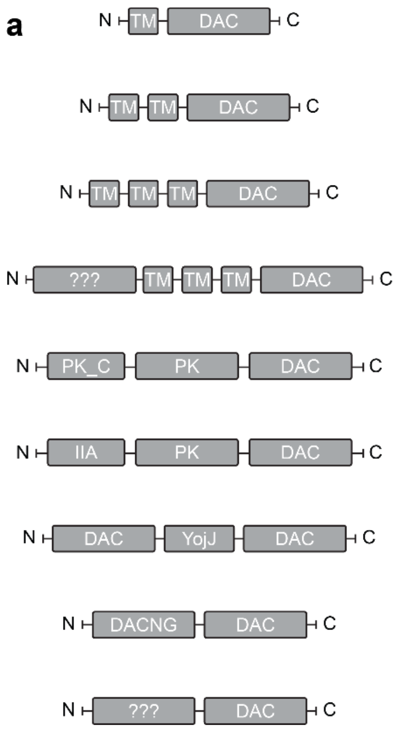


**Extended Data Fig. 1: Our search identified novel non-immune DACs with unrecognized domain organizations.**

a) A small subset of the non-immune DACs found in this study which had not been identified in the literature.

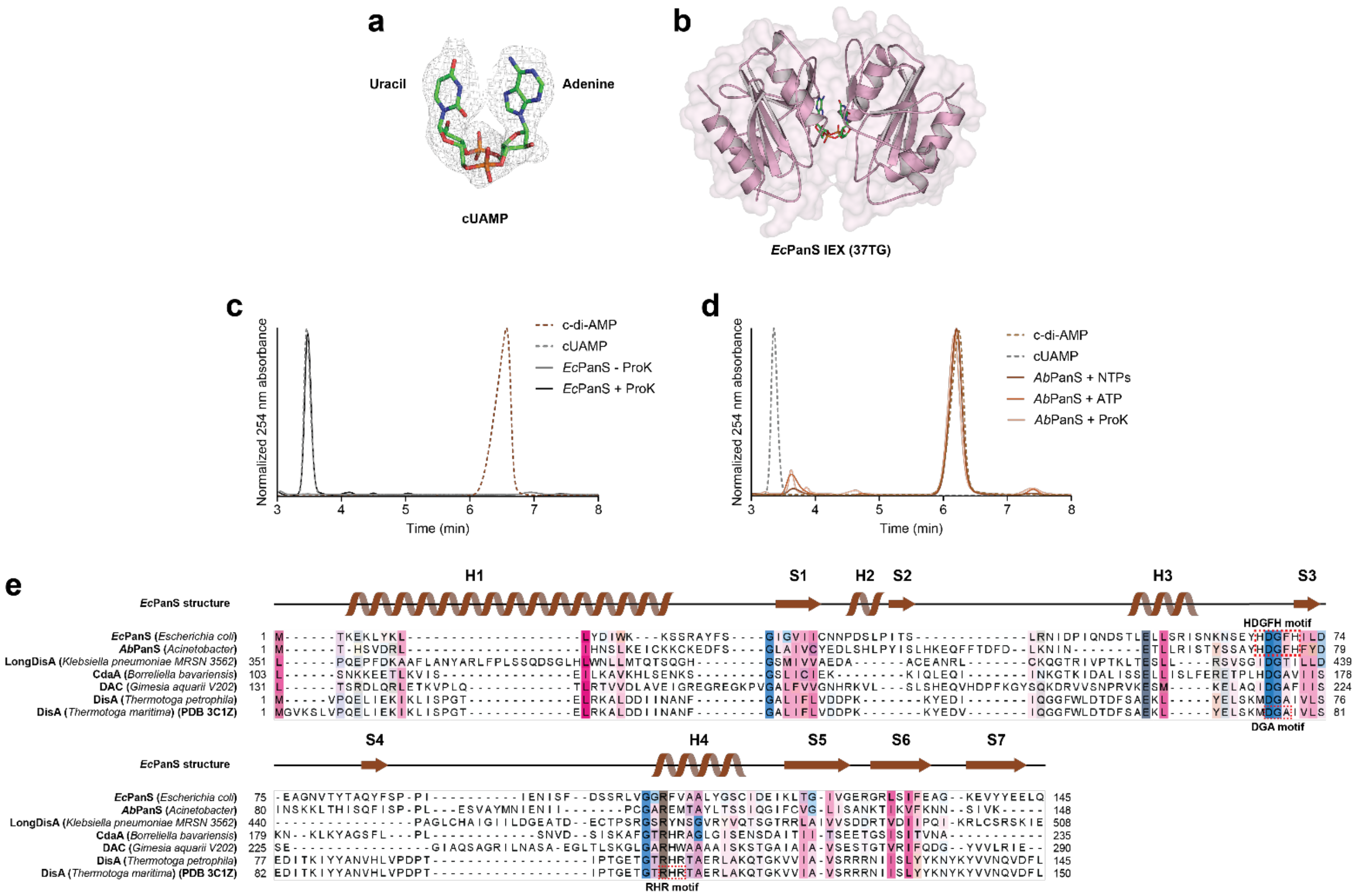


**Extended Data Fig. 2: *Ec*PanS copurifies with cUAMP while other homologs do not.**

a) Polder map of cUAMP from the non-anion-exchanged *Ec*PanS structure (37TH)

b) Crystal structure of anion-exchanged *Ec*PanS bound to cUAMP (green).

c) High-performance liquid chromatography data for the Proteinase K digestion of *Ec*PanS showing the release of cUAMP. Representative of three biological replicates.

d) High-performance liquid chromatography data for enzymatic reactions of *Ab*PanS with NTPs or with ATP. Representative of three biological replicates.

e) Alignment of several DACs identified in this study as well as the *Tm*DisA mentioned throughout this work (bottom). The secondary structure of *Ec*PanS is shown in the top row for comparison.

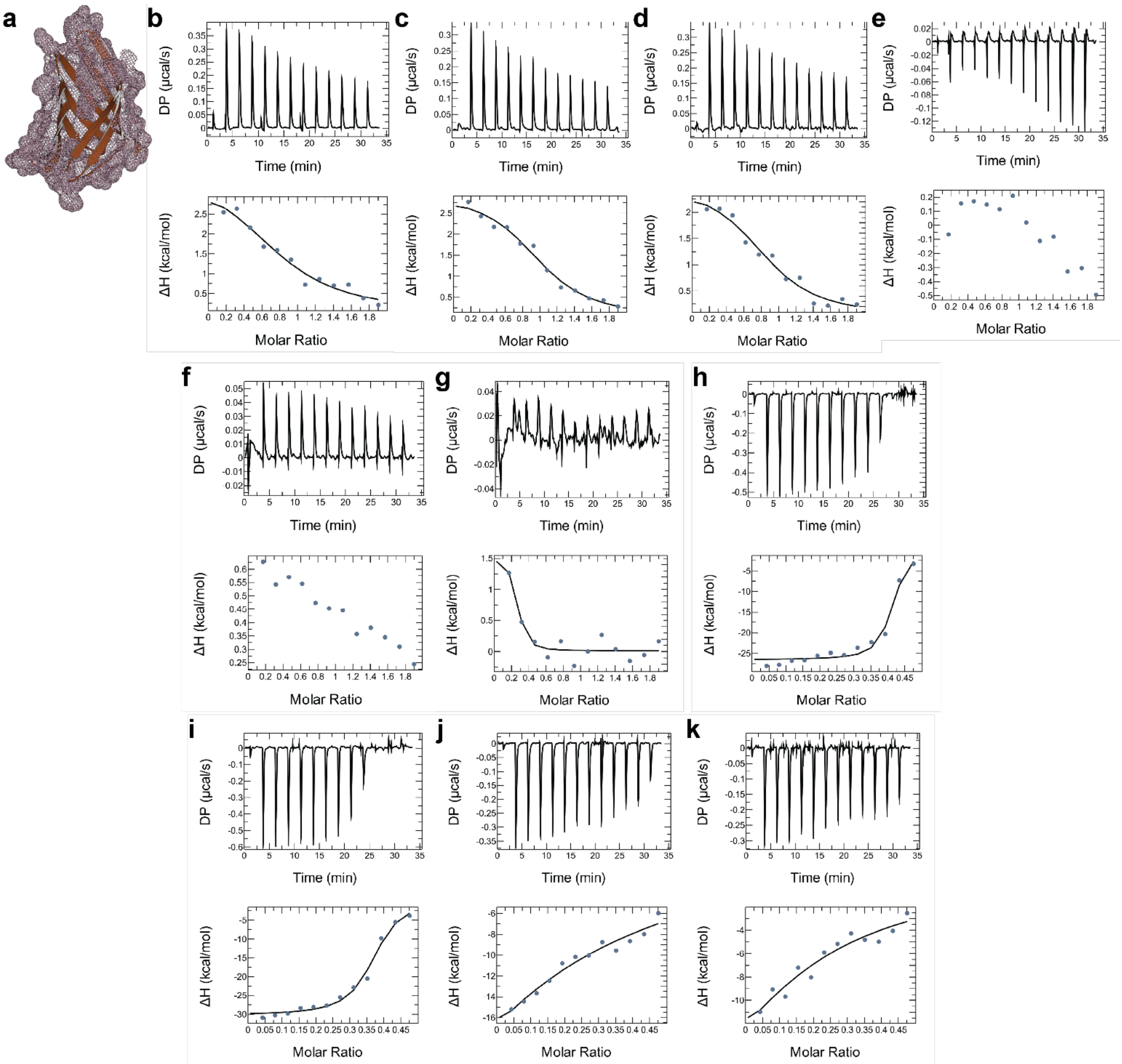


**Extended Data Fig. 3: ITC data for c-di-AMP and cUAMP binding to *Ec*PanE and N4 Acb4.**

a) Surface of *Ec*PanE AF3 structure showing predicted c-di-AMP binding pocket.

b-d) Raw ITC readout for *Ec*PanE binding to c-di-AMP

e-g) Raw ITC readout for *Ec*PanE binding to cUAMP

h-i) Raw ITC readout for N4 Acb4 binding to c-di-AMP

j-k) Raw ITC readout for N4 Acb4 binding to cUAMP
